# Natural selection on *TMPRSS6* associated with the blunted erythropoiesis and improved blood viscosity in Tibetan Pigs

**DOI:** 10.1101/380543

**Authors:** Xiaoyan Kong, Xinxing Dong, Shuli Yang, Jinhua Qian, Jianfa Yang, Qiang Jiang, Xingrun Li, Bo Wang, Dawei Yan, Shaoxiong Lu, Huaming Mao, Xiao Gou

**Affiliations:** Faculty of Animal Science and Technology, Yunnan Agricultural University, Kunming, Yunnan, China; Department of Animal Science, Yuxi Agriculture Vocational-Technical College, Yuxi, Yunnan, China; Dairy Cattle Research Center, Shandong Academy of Agricultural Science, Jinan, Shandong, China; Department of Animal Science, Dali Vocational and Technical College of Agriculture and Forestry, Dali, Yunnan, China; Research Experimental Center, Yunnan University of Traditional Chinese Medicine, Kunming, Yunnan, China

**Author notes:** Corresponding authors (HMM); (XG). These authors contributed equally to this work (Xiaoyan Kong, Xinxing Dong, Shuli Yang).

**Keywords:** Tibetan Pigs, *TMPRSS6*, HGB, CGTG indel, High-altitude adaptation

## Abstract

Tibetan pigs, indigenous to Tibetan plateau, are well adapted to hypoxia. So far, there have been not any definitively described genes and functional sites responsible for hypoxia adaptation for the Tibetan pig. Here we conducted resequencing of the nearly entire genomic region (40.1 kb) of the candidate gene *TMPRSS6* (Transmembrane protease, serine 6) associated with hemoglobin concentration (HGB) and red blood cell count (RBC) in 40 domestic pigs and 40 wild boars from five altitudes along the Tea-horse ancient road and identified 708 SNPs, in addition to an indel (CGTG/----) in the intron 10. Both the CGTG deletion frequency and the pairwise r^2^ linkage disequilibrium showed an increase with elevated altitudes in 838 domestic pigs from five altitudes, suggesting that *TMPRSS6* has been under Darwinian positive selection. As the conserved core sequence of hypoxia-response elements (HREs), the deletion of CGTG in Tibetan pigs decreased the expression levels of *TMPRSS6* mRNA and protein in the liver revealed by real-time quantitative PCR and western blot, respectively. To explore whether reduced *TMPRSS6* expression level could improve blood viscosity, the relationship between CGTG indel and hematologic and hemorheologic parameters in 482 domestic pigs from continuous altitudes was detected and dissected a genetic effect on reducing HGB by 13.25g/L in Gongbo’gyamda Tibetan pigs and decreasing MCV by 4.79 fl in Diqing Tibetan pigs. In conclusion, the CGTG deletion of *TMPRSS6* resulted in lower HGB and smaller MCV, thereby blunting erythropoiesis and improving blood viscosity as well as erythrocyte deformability.

## Introduction

The Tibetan pig lives at an average elevation of approximately 3500 m on Tibetan plateau [1], due to the availability of well-adapted extreme conditions including low ambient temperature, high ultraviolet radiation, harsh climate, and low oxygen [2]. For thousands of years, Tibetan pigs have developed complex phenotypic and physiological adaptations to high-altitude hypoxia compared with lowland pigs [3]. When animals are exposed to chronic hypoxia, the pulmonary artery pressure and red blood cell count (RBC) increase, causing pulmonary hypertension and right ventricular hypertrophy. Experimental evidence indicates that pulmonary artery pressure, pulmonary artery wedge pressure, cardiac output and pulmonary vascular resistance increase with increased elevations in pigs [4].

The hypoxia-induced increase in RBC and HGB, thereby rising blood viscosity and resistance, result in pulmonary hypertension and right-heart failure [5]. In the past few years, increasing attentions have been devoted to identify the genes related to blood physiology underlying the adaptation to high-altitude hypoxia for Tibetans [6–9], yaks [10, 11], Tibetan chickens [12], Tibetan antelopes [13], pikas [14], Tibetan pigs [2, 3, 15], and deer mice [16]. Multiple genome-wide scans showed that positive selection in human beings and animals at high altitude occurred mainly in the hypoxia-inducible factor (HIF) signaling pathway [3, 6, 17–24]. In this pathway, *EPAS1* (endothelial PAS domain protein 1) and *EGLN1* (egl nine homolog 1) are key genes that correlated significantly with hemoglobin concentration [6, 11, 23, 25–27]. The genome-wide associated with blood physiology also revealed that the gene *TMPRSS6* (Transmembrane protease, serine 6, one of the downstream genes in the HIF pathway) was strongly correlated with serum iron concentration [28, 29, 38–43, 30–37], RBC, and HGB [44–46]. This gene encodes a type II transmembrane serine proteinase (Matriptase 2) expressing in the liver [47], and regulates the iron metabolism by controlling the hepcidin expression [33, 35, 37, 39, 48, 49]. A genome-wide meta-analysis also identifies *TMPRSS6* associated with hematological parameters in the HaemGen consortium [46]. Current evidence allso suggests that the *TMPRSS6* variants were significantly associated with plasma ferritin, hemoglobin, risk of iron overload, and type 2 diabetes in Chinese Hans [50]. Several lines of evidence, mutations in *TMPRSS6* are known to be associated with cause iron-refractory iron deficiency anemia [36, 37, 41, 42, 51, 52] or iron deficiency[31, 35, 53–55], especially one mutation 736 (V) allele (rs855791) allele tended to be associated with low Hb levels and iron status in humans [35, 36, 38, 39, 44, 45, 56].

In this study, we aim to probe into the potential role of the *TMPRSS6* gene underlying blood physiological adaptation to high-altitude Tibetan pigs. Resequencing of the nearly complete genomic region of *TMPRSS6* (40.1 kb) was conducted to explore the molecular mechanism of adaptation to high-altitude hypoxia in Tibetan pigs. Otherwise, real-time quantitative PCR and western blot were performed to determine the expression level of *TMPRSS6*. Meanwhile, hematologic and hemorheologic parameters were measured to analyze the association with *TMPRSS6*, including red blood cell count (RBC), hemoglobin concentration (HGB), hematocrit (HCT), mean corpuscular volume (MCV), mean corpuscular hemoglobin (MCH), mean corpuscular hemoglobin concentration (MCHC) and red blood cell distribution width (RDW).

## Materials and methods

### Ethics Statement

All animal experimental procedures were performed according to the Guide for the Care and Use of Laboratory Animals (Ministry of Science and Technology of China, 2006).

### Animal and sample preparation

In this study, a total of 838 domestic pigs were included, in which 353 Tibetan pigs were from 5 different locations in Tibetan plateau and 485 pigs were from 9 domestic breeds across the other three altitudes along the Tea-horse ancient road, and 40 Chinese wild boars were sampled from 5 continuous altitudes along the Tea-horse ancient road. Sample size and localization of each population were shown in Supplementary Table S1 and Fig 1. 80 individuals including 40 domestic pigs and 40 Chinese wild boars were used for *TMPRSS6* resequencing. 838 pigs were genotyped for haplotype analyze of *TMPRSS6* gene. Hematologic and hemorheologic parameters (Table S2) of 482 pigs from the 838 domestic pigs were detected. Ten Gongbo’gyamda Tibetan pigs with genotype DD (n = 5) and II (n = 5) were selected to detect expression level of mRNA and protein by real-time quantitative PCR (RT-qPCR) and western-blot experiments. Liver tissue specimens were collected and immediately frozen in liquid nitrogen.

**Fig 1.**
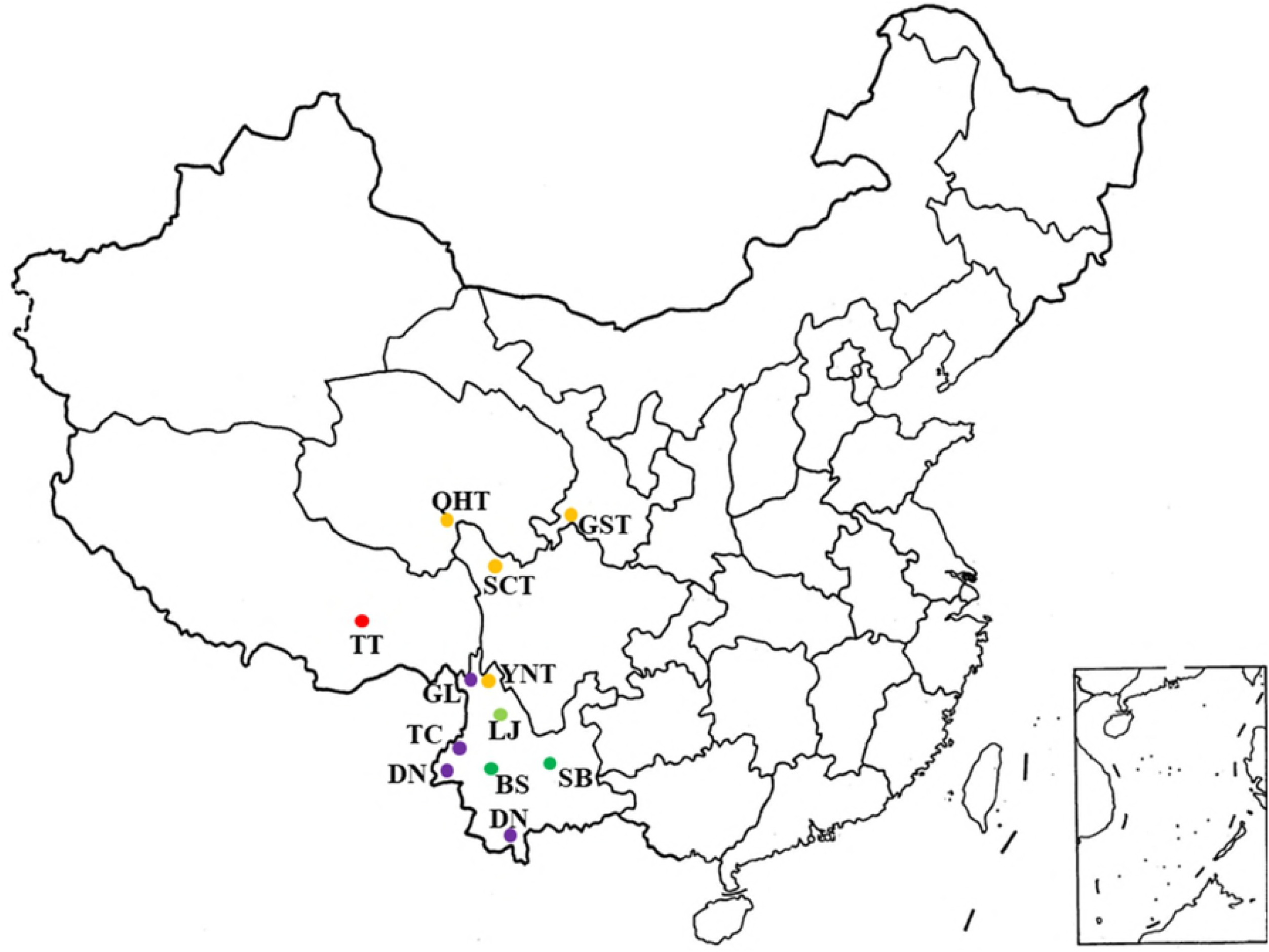
Sampling and geographic locations of the pig breeds. The red, yellow, green, blue and purple indicated high level of high altitude (HHA, 3600 meters), middle level of high altitude (MHA, 3300 meters), low level of high altitude (LHA, 2500 meters), middle level of altitude (MH, 1600 meters) and low level of altitude (LA, 550 meters), respectively. TT, Tibetan (Gongbo’gyamda); YNT, Tibetan (Yunnan); SCT, Tibetan (Sichuan); QHT, Tibetan (Qinghai); GST, Tibetan (Gansu); LJ, Lijiang; BS, Baoshan; SB, Saba; DN, Diannanxiaoer; TC, Tengchong; GL, HighLiGongshan.

### Resequencing, genotyping and haplotype analysis

The *TMPRSS6* gene (Genbank accession no. NC_010447) was resequenced in 40 domestic pigs and 40 wild boars across 5 continuous altitudes, 10 individuals per altitude. A sample of 5 mL blood from each individual pig was obtained from the jugular vein. Blood genomic DNA was extracted with a Genomic DNA Isolation Kit (Bactec, Beijing, China) according to the manufacturer’s instructions. We designed 55 pairs of primers to amplify the entire gene, including 17 pairs of primers for exons and 38 pairs of primers for introns (Table S3). PCR was performed in 25 μL reaction volume. The ingredient comprised 2 μL DNA template, 11μL Mix, 1 μmol/L of each forward and reverse primer and 10 μL ddH_2_O. PCR procedure was performed as following: initial denaturation for 5 min at 95 °C, followed by 36 cycles of 95°C for 30 s; annealing at prescribed annealing temperature (Table S3) for 30 s; and extension at 72°C for 45 s. The final extension was performed at 72°C for 8 min. PCR products were sequenced using an ABI 3730 sequence analyzer (Applied Biosystems, Foster City, CA). *TMPRSS6* SNPs were genotyped also by Sanger sequencing. The sequence polymorphisms were analyzed using MEGA7 software[57]. The LD map of *TMPRSS6* SNPs in the domestic pigs and wild boars was constructed by Haploview using the r^2^ algorithm [58].

### Hematologic and Hemorheologic parameters

Hematologic parameters were measured by BC-2800Vet Auto Hematology Analyzer (Mindray Co., Ltd.). The following measurements were obtained: red blood cell count (RBC), hemoglobin concentration (HGB), hematocrit (HCT), mean corpuscular volume (MCV), mean corpuscular hemoglobin (MCH), mean corpuscular hemoglobin concentration (MCHC) and red blood cell distribution width (RDW), respectively. Hemorheologic parameters were measured using a ZL1000 Auto Blood Rheology Analyzer (Zonci Co., Ltd.) at a constant temperature of 37°C. The following measurements were obtained: fibrinogen (FB), plasma viscosity (PV), whole blood relative index of low shear (WLS) at shear rate of 1 S^−1^, whole blood relative index of middle shear (WMS) at shear rate of 5 S^−1^, whole blood relative index of high shear (WHS) at shear rate of 200 S^−1^, RBC aggregation index (EAI), RBC aggregation coefficient (EAC), casson viscosity (CV), RBC internal viscosity (RBCIV), low shear flow resistance (LSFR) at shear rate of 1 S^−1^, middle shear flow resistance (MSFR) at shear rate of 5 S^−1^, high shear flow resistance (HSFR) at shear rate of 200 S^−1^, yield stress (YS) respectively. Parameters were measured immediately after veinpuncture.

### Real-time quantitative PCR

Total RNA was extracted from the livers with total RNA extraction kit(DP419) (Tiangen, Beijing, China). The concentration and purity of RNA were determined with a NanoDrop 2000 Biophotometer (Thermo Fisher Scientific Inc., West Palm Beach, FL, USA) and integrity was verified by electrophoresis in a 1% agarose gel. After treatment with DNase I, 2 μg of RNA in a 20 μL reaction volume was reversely transcribed into cDNA using a cDNA Kit (TaKaRa, Dali, China). To avoid genomic DNA contamination, Primer Premier 5.0 software was used to design *TMPRSS6* gene (XM_001924749) primers that amplified products spanning an intron. The primers were 5′-CCCGATTCGCTCTTCTCC-3′ and 5′-GGCACCTTCCTTTCA CCC-3′. The 28s rRNA (DQ297674) was used as the internal standard and its primers were 5′-CGGGATGAACCGAACGC-3′ and 5′-GCCACCGTCCTGCTGTCT-3′. Real-time quantitative PCR (RT-qPCR) was conducted on the Bio-Rad CFX96 System (Bio-Rad, USA). Each reaction mixture contained 10.0 μL 2×SYBR Green qPCR SuperMix (Transgen, Beijing, China), 1.0 μL cDNA, 0.5 μL of each primer (10.0 nmol/μL), ddH_2_O water was added to adjust the volume to 20.0 μL. The RT-qPCR program started with denaturation at 95 °C for 15 min followed by 39 cycles of denaturation at 95 °C for 10 s and annealing elongation at 63 °C for 30 s, during which fluorescence was measured. Next, a melting curve was constructed by increasing the temperature from 65 °C to 95 °C in sequential steps of 0.5 °C for 5 s, during which fluorescence was measured. The RT-qPCR efficiency of each pair of primers was calculated using 5 points in a 5-fold dilution series of cDNA, which was used to construct a standard curve. A cDNA pool of all samples was used as a calibration and three replications of each sample were performed. Gene expression levels were calculated using the 2^−ΔΔCt^ method (ΔΔCt = ΔCt target gene - ΔCt 28srRNA gene) as previously described [59].

### Western blot

The liver tissue was homogenized using a Mixer MillMM400 (Retsch, Germany) for 5 min and then centrifuged at 10,000 × g for 10 min at 4 °C. Protein concentrations were determined using a Protein Assay Kit (Bio-Rad). Proteins (40 μg) were separated by sodium dodecyl sulfate polyacrylamide gel electrophoresis (SDS-PAGE) using a 5% stacking gel and a 10 % separating gel. Following electrophoresis, proteins were transferred to Immobilon-P Transfer Membranes (IPVH00010) for 2 h at 300 mA using a Bio-Rad Criterion Blotter. Membranes were blocked overnight in blocking buffer (P0023B, Beyotime Ltd., China) and then incubated with primary mouse monoclonal *GAPDH* (1:1,000 dilution, AG019, Beyotime Ltd., China) and Anti-tmpss6 (n-terminal) polyclonal antibody (1:500 dilution, Sigma) diluted in primary antibody dilution buffer (P0023A, Beyotime Ltd., China) at 4 °C for 2 h. The membranes were washed 3 times with PBST (phosphate buffer saline containing 0.1 % Tween 20), and incubated with secondary goat anti-rabbit (1:1,000 dilution, A0216, Beyotime Ltd., China) antibody diluted in secondary antibody dilution buffer (P0023D, Beyotime Ltd., China) for 1 h. After the membranes were washed 3 times in Tris-buffered saline with Tween for 30 min, immune complexes were visualized using an eECL Western Blot Kit (CW0049A, CWBIO Ltd., China) according to the manufacturer’s instructions. To determine expression ratio of TMPRSS6, western blot was analyzed using Image J 1.44 software (NIH, USA).

### Statistical analysis

Correlation coefficient between the altitudes and frequencies of CGTG deletion were calculated using the *F* test in SAS software (ver. 9.0). Candidate gene association of *TMPRSS6* SNP genotypes with Hematologic and Hemorheologic parameters was performed only within the same population or the population at the same elevation. The *F* test in the analyze of variance (ANOVA) in the SAS software (ver. 9.0) was used to analyze the association based on the following linear model: Y_ijkl_=μ + G_i_ + S_k_ + Aj + e_ijkl_. Where, Y_ijkl_ was the phenotypic value of the target trait, μ was the population mean, G_i_ was the effect of the i^th^ genotype, S_k_ was the effect of k^th^ gender, Aj was the effect of the j^th^ age, and e_ijkl_ was the random residual [60].

## Results

### Complete sequence of *TMPRSS6* and discovery of a selected CGTG indel

To reveal the detailed pattern of sequence variations of *TMPRSS6* in pigs, resequencing of the entire genomic region of *TMPRSS6* (about 40.1 kb) was conducted firstly covering 30.1-kb exon–intron region, 7.4-kb 5’-region and 0.6-kb 3’-region. Totally, 80 pigs were resequenced including 40 domestic pigs and 40 wild boars across continuous altitudes along the ancient Tea-horse road. A total of 708 SNPs (Table S1) were screened and all of variations were synonymous SNPs. The linkage disequilibrium (LD) plot of 708 SNPs in 4 domestic pig populations and 4 wild boar population were shown in Fig 2. Tibetan pigs and Tibetan wild boars living in high altitude above 3000 m displayed higher LD compared with the other pigs. And the LD increased with elevated altitudes, suggesting that altitude selection on *TMPRSS6* has been consistent in both domestic pigs and wild boars. Specifically, an CGTG indel occurred in the intron 10 of *TMPRSS6* gene, closely linked with SNP13, SNP15 and SNP16. The CGTG acted as the conserved core sequence of hypoxia-response elements (HREs) [61] which bound to many oxygen-regulated genes by hypoxia induced factors (HIFs). Therefore, extensive samples were needed to determine the intron 10 sequence of *TMPRSS6* gene to further reveal whether the CGTG indel played a potential role in high-altitude adaptation of Tibetan pigs.

**Fig 2.**
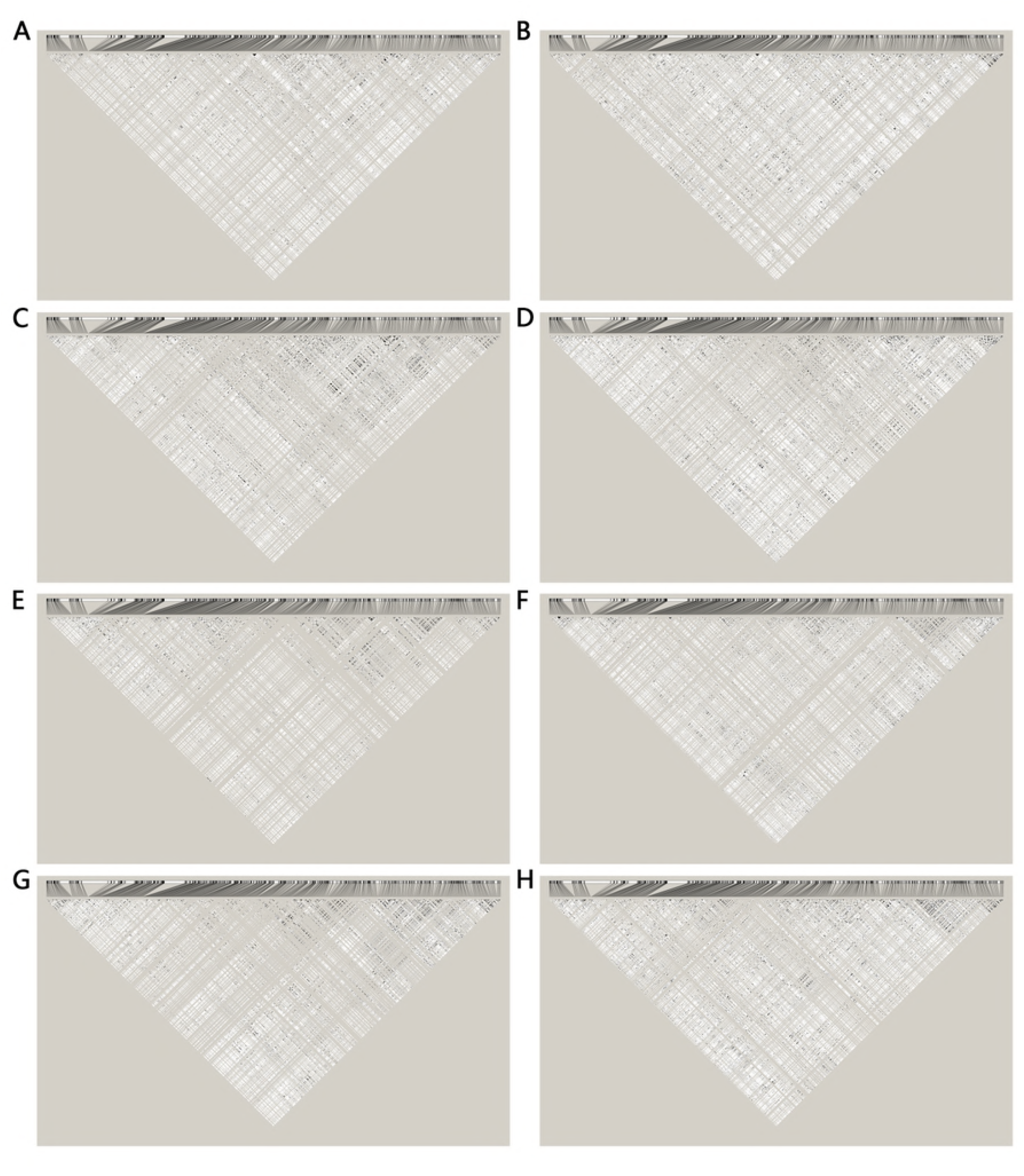
Pairwise r^2^ linkage disequilibrium spanning entire *TMPRSS6* gene in pig breeds along continuous altitudes. A total of 708 SNPs of nearly whole TMPRSS6 gene from 40 domestic pigs and 40 wild boars were included. The SNPs with minor allele frequency smaller than 10% were removed. Darker shading indicates higher levels of LD. (A) LA domestic pigs; (B) LA wild boars; (C), MA domestic pigs; (D) MA wild boars; (E) LHA domestic pigs; (F) LHA wild boars; (G) MHA domestic pigs; (H) MHA wild boars.

### Positive selection analysis of intron 10 in *TMPRSS6* gene

To test whether the observed CGTG indel was a hypoxia-selected site, the intron 10 of *TMPRSS6* gene was sequenced in a total of 838 domestic pigs from 5 altitudes along the Tea-horse ancient road (Table S1 and Fig 1). As a result, 16 SNPs and the CGTG indel were screened. Genotype frequency and allele frequency were listed in Table S4. In the 5 altitude pig populations from 500 m to 3500 m, the frequencies of CGTG deletion in domestic pigs were 44.3%, 46.9%, 51.8%, 53.7% and 62.2%, respectively. In addition, the correlation between the allele frequency of CGTG indel and altitude was analyzed and a strong positive correlation (r=0.959, P=0.005) in domestic pigs was found (Fig 3 and Fig 4), implicating that the prevalent CGTG deletion might play a key role in contributing to high-altitude hypoxia adaptation in Tibetan pigs. Pairwise r^2^ linkage disequilibrium for the 16 SNPs were separately analyzed in pig breeds across continuous altitudes, and elevated r^2^ values among the CGTG indel and its three neighboring sites (SNP13, SNP15, SNP16) with increased altitudes (Fig 5) suggested a genetic hitchhiking effect in domestic pigs across continuous altitudes.

**Fig 3.**
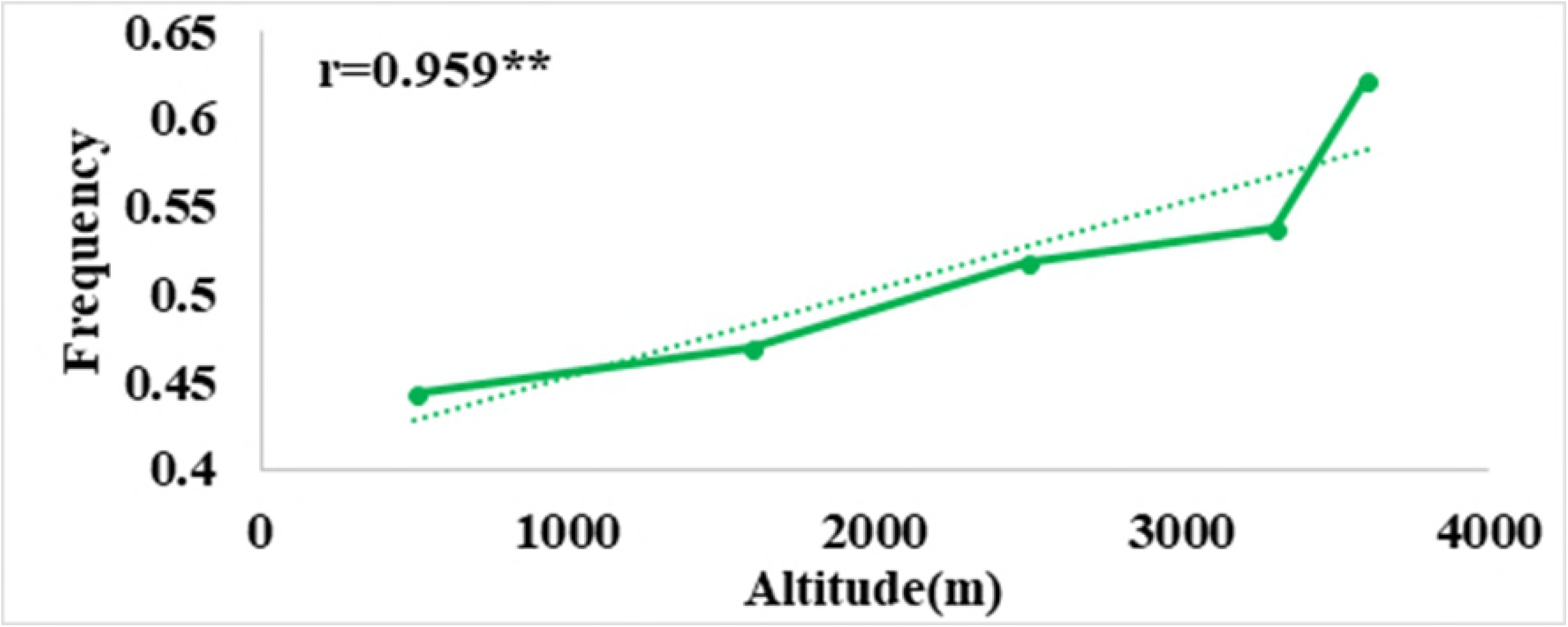
The correlation between CGTG deletion frequency and altitude in domestic pigs. Plot of the correlation analysis between frequencies of the CGTG deletion and the altitudes among domestic pigs, “r” represents the correlation coefficient (r = 0.959, P = 0.005).

**Fig 4.**
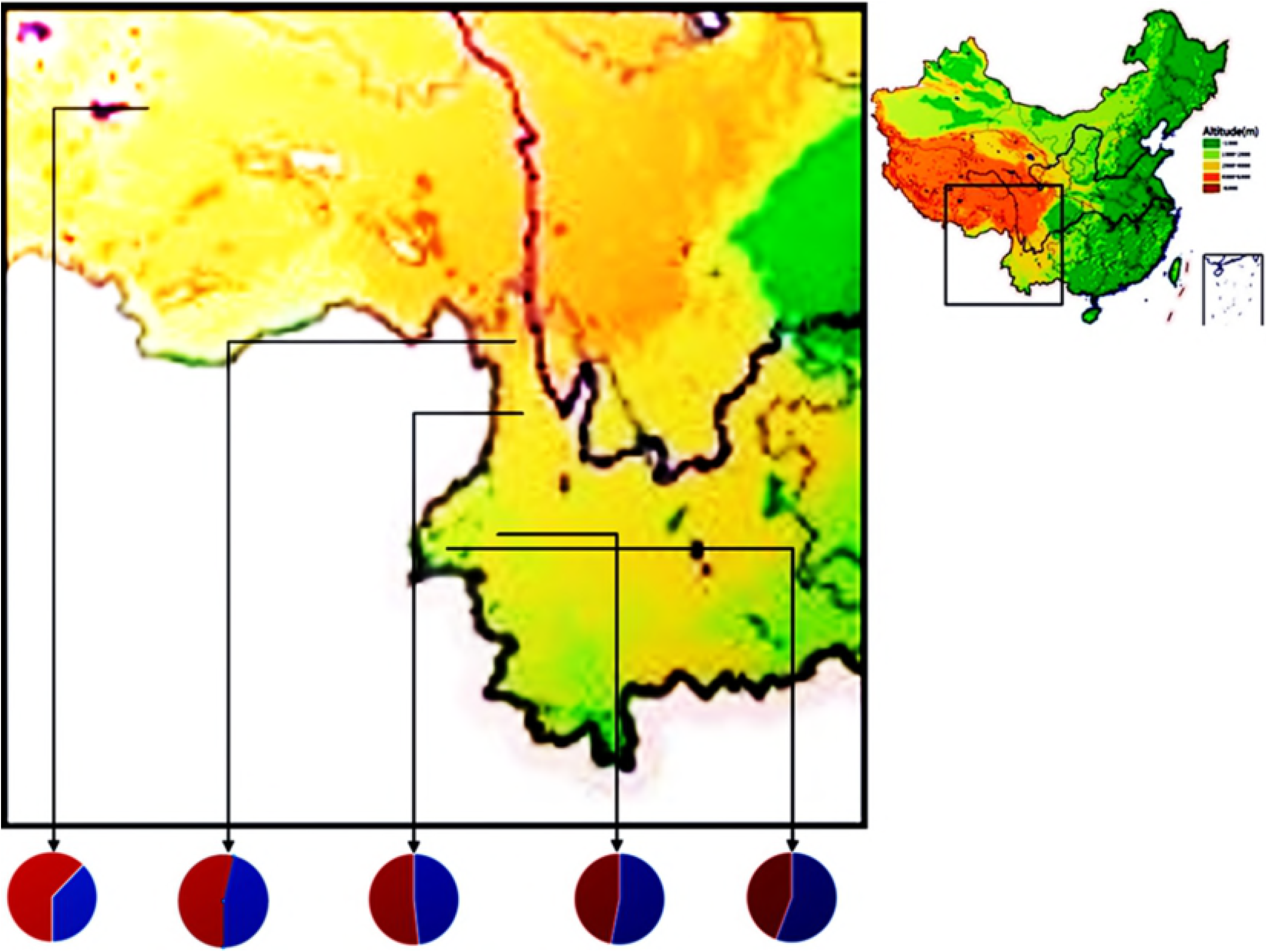
Altitudinal pattern of allele frequency variation at the CGTG indel in pig breeds across continuous altitudes. In the pie diagram, the frequency of the deletion allele was shown in red. From south to north, the sampling altitudes were HHA (3500 m), MHA (3000 m), LHA (2500 m), MA (1500 m) and LA (500 m).

**Fig 5.**
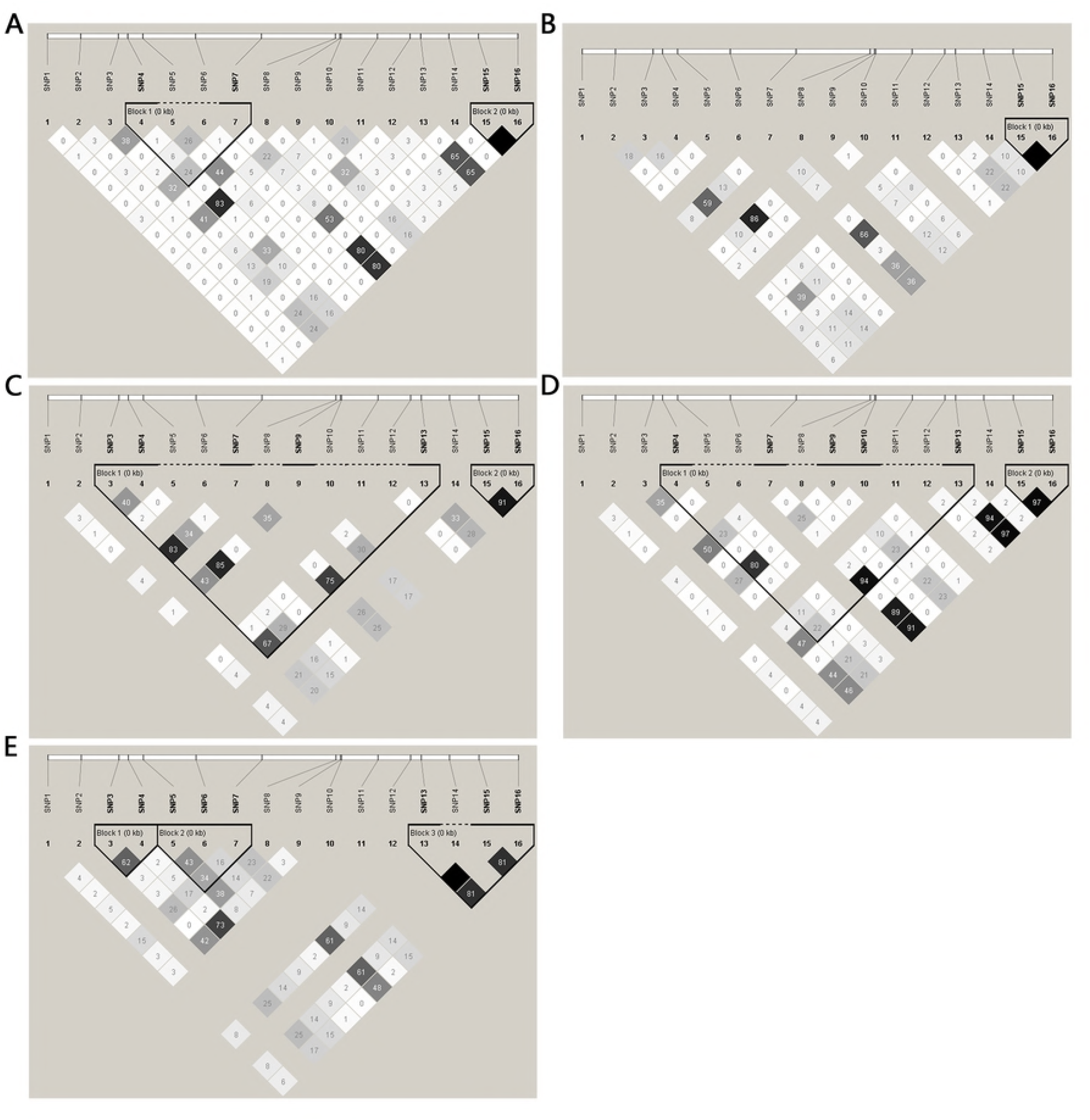
Pairwise r^2^ linkage disequilibrium spanning the intron 10 in pig breeds across continuous altitudes. A total of 16 SNPs from 838 domestic pigs were included. The SNPs with minor allele frequency smaller than 10% were removed. Darker shading indicated higher levels of LD. (A) LA domestic pigs; (B) MA domestic pigs; (C) LHA domestic pigs; (D) MHA Tibetan pigs; (E) HHA Tibetan pigs.

### Association of the CGTG indel with blood physiological parameters

Genome-wide association study had established the association between variants in *TMPRSS6* and hemoglobin level in humans [44, 46]. To further investigate the role of the CGTG deletion of *TMPRSS6* in blood physiological adaptation, we genotyped the CGTG indel and measured 20 indexes of the hematologic and hemorheologic parameters in two high-altitude Tibetan pig populations (HHA and MHA) and three other altitude pig populations (LHA, MH and LA). As shown in Table 1, the CGTG indel was significantly associated with HGB and RBC in HHA Tibetan pigs, and both RBC and HGB value of the insertion homozygote (II) were significantly higher than those of the deletion homozygote (DD) (p<0.01), which was accompanied by similar statistically tendencies for hemorheologic parameter WLS, WMS and WHS (p<0.05). The sex- and age-adjusted HGB and RBC were 13.25 g/L and 0.64×10^6^/ul lower in the DD genotype compared with the II genotype in HHA Tibetan pigs. By contrast, the significant differences between genotypes for hematologic parameter HGB, MCV, and hemorheologic parameter CV, YS, LSFR, MSFR, HSFR (p<0.01 for MCV, p<0.05 for the others) occurred in MHA Tibetan pigs (Table 2). There were no significant differences for hematologic and hemorheologic parameters in the other three altitude breeds (p>0.05) (S5-7 Tables), suggesting that the CGTG deletion might play important roles in lowering HGB to improve blood viscosity.

**Table 1.**
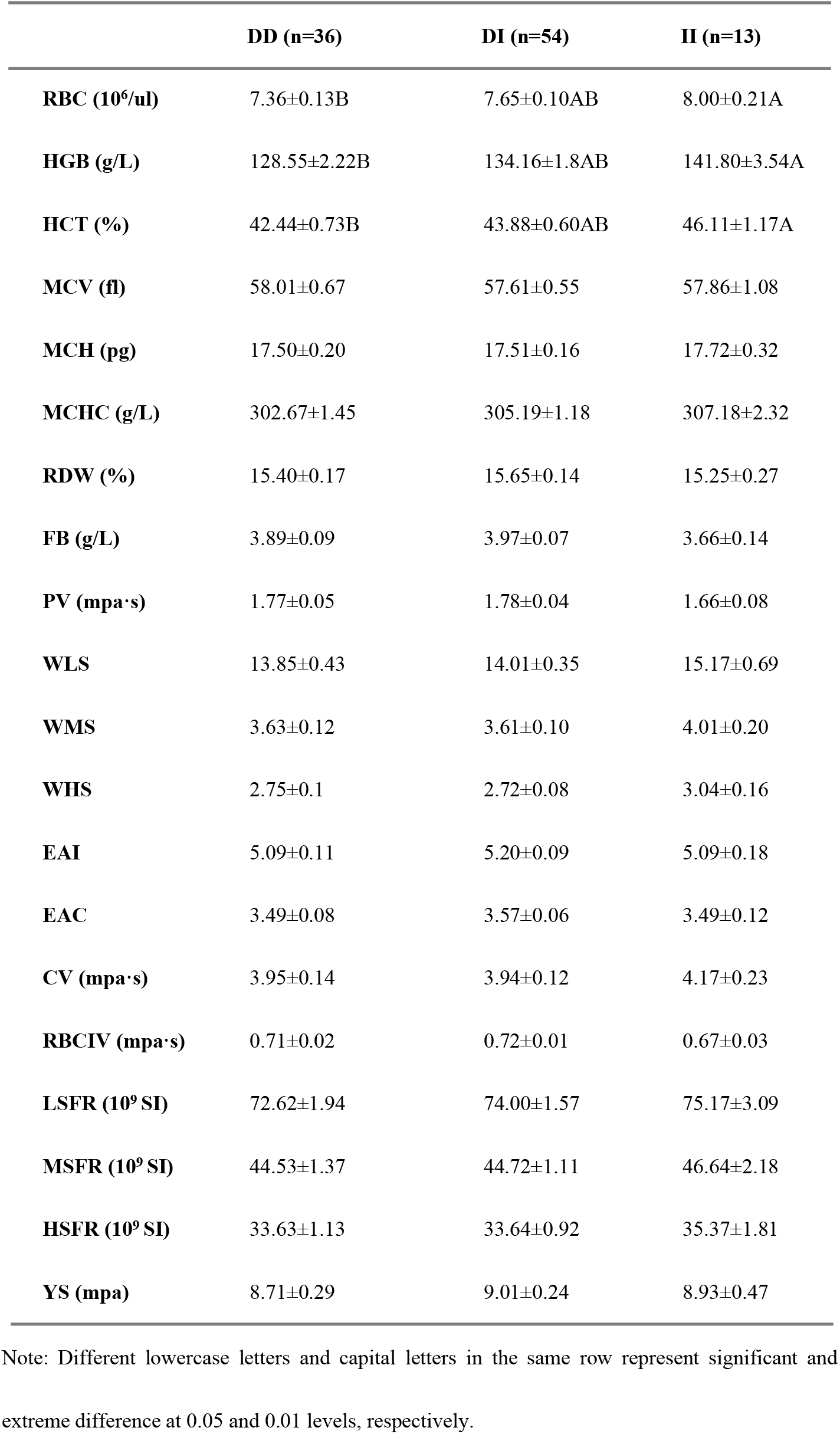
Association of the CGTG indel with hematologic and hemorheologic parameters in Tibetan pigs at 3600 m.

**Table 2.** Association between CGTG indel and hematologic and hemorheologic parameters in Tibetan pigs at 3300 m.

|  | **DD (n=12)** | **DI (n=17)** | **II (n=15)** |
| --- | --- | --- | --- |
| **RBC (10<sup>6</sup>/ul)** | 7.19±0.25 | 7.49±0.20 | 7.08±0.19 |
| **HGB (g/L)** | 127.07±3.65b | 134.80±2.82ab | 136.65±2.96a |
| **HCT (%)** | 41.03±1.09b | 43.58±0.84a | 44.60±0.88a |
| **MCV (fl)** | 57.30±1.38B | 60.08±1.12AB | 62.09±1.07A |
| **MCH (pg)** | 17.67±0.46b | 18.31±0.38a | 19.14±0.36a |
| **MCHC (g/L)** | 309.20±2.75 | 305.86±2.23 | 308.56±2.13 |
| **RDW (%)** | 16.00±0.34 | 15.42±0.28 | 14.92±0.27 |
| **FB (g/L)** | 4.81±0.34 | 5.43±0.28 | 5.22±0.26 |
| **PV (mpa·s)** | 2.17±0.16 | 2.47±0.13 | 2.37±0.12 |
| **WLS** | 9.75±0.78 | 9.12±0.64 | 10±0.61 |
| **WMS** | 2.53±0.19 | 2.38±0.15 | 2.57±0.15 |
| **WHS** | 1.89±0.14 | 1.78±0.11 | 1.91±0.11 |
| **EAI** | 5.17±0.16 | 5.06±0.13 | 5.23±0.12 |
| **EAC** | 3.53±0.11 | 3.47±0.09 | 3.6±0.08 |
| **CV (mpa·s)** | 3.18±0.13 | 3.48±0.11 | 3.57±0.10 |
| **RBCIV (mpa·s)** | 0.89±0.06 | 0.98±0.05 | 0.95±0.05 |
| **LSFR (10<sup>9</sup> SI)** | 60.10±3.60 | 64.87±2.92 | 68.56±2.79 |
| **MSFR (10<sup>9</sup> SI)** | 36.26±1.65 | 39.51±1.34 | 40.87±1.28 |
| **HSFR (10<sup>9</sup> SI)** | 27.24±1.18 | 29.78±0.95 | 30.62±0.91 |
| **YS (mpa)** | 7.32±0.59 | 7.82±0.48 | 8.48±0.45 |

Note: Different lowercase letters and capital letters in the same row represent significant and extreme difference at 0.05 and 0.01 levels, respectively.

### *TMPRSS6* mRNA and protein expression

To investigate the regulatory effect of CGTG deletion, *TMPRSS6* mRNA and protein expression level in livers [62] of HHA Tibetan pigs with genotype DD and II were detected, respectively. As shown in Fig 6A, the expression level of *TMPRSS6* mRNA in liver was significantly lower in DD genotype (n = 5) compared with that in the II genotype (n = 5) in HHA Tibetan pigs (P < 0.01). In addition, the matriptase-2 protein expression level was strikingly lower in DD genotype (n = 5) than that of the II genotype (n = 5) in HHA Tibetan pigs (Figs 6B and 6C). The consistent results from RT-qPCR and western blot showed that the CGTG deletion leaded to the lower expression level of mRNA and protein in *TMPRSS6*.

**Fig 6.**
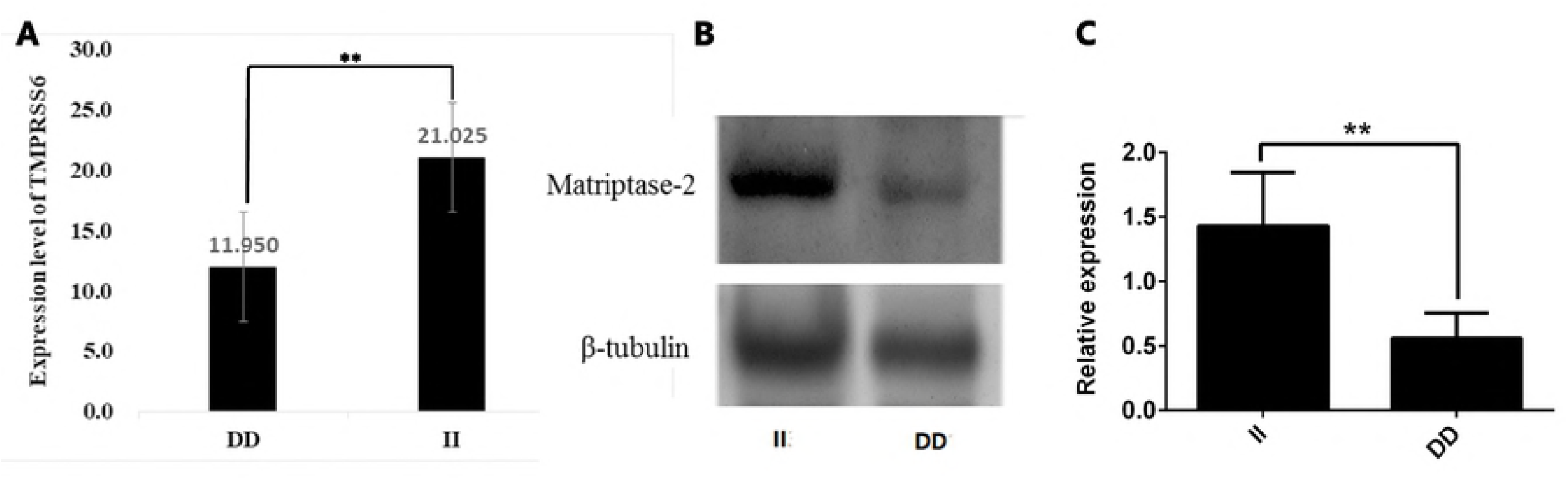
mRNA and protein expression levels of *TMPRSS6* in the livers of HHA Tibetan pigs. (A) The mRNA level of *TMPRSS6* in the livers of HHA Tibetan pigs with the DD (n = 5) and II (n = 5) genotype was measured by RT-qPCR. Error bars represent mean ± SD from five independent experiments. **P < 0.01. (B) and (C) Western Blot was performed to detect the expression level of matriptase-2 protein encoded by *TMPRSS6* in HHA Tibetan pigs with the DD and II genotype. β-tubulin expression was used as the loading control. DD, deletion (n = 3); II, insertion (n = 3).

## Discussion

The Tibetan pig is one of typical high altitude domestic animals. *TMPRSS6* is the typical gene correlated with the high-altitude adaption of the Tibetan pigs. And the protein encoded by *TMPRSS6* is Matriptase 2 that expresses in the liver and regulates the expression of the systemic iron regulatory hormone hepcidin [51]. The hepatic hormone hepcidin regulates body iron metabolism based on the two mechanisms, the “stores regulator” and the “erythroid regulator” [63–69]. Genome-wide association studies (GWAS) have found that four erythrocyte traits, including Hb, Hct, MCH and MCV, were significantly associated with *TMPRSS6* in human [38, 44, 70]. To explore the molecular mechanism of adaptation to high-altitude hypoxia in Tibetan pigs, we conducted resequencing of the nearly complete genomic region of *TMPRSS6* (40.1 kb) in 40 domestic pigs and 40 wild boars and identified one CGTG deletion highly prevalent in Tibetan pigs. Haplotype analysis of sequence variations in *TMPRSS6* gene in pig populations along continuous altitudes revealed a hitchhiking effect close to CGTG deletion, suggesting that *TMPRSS6* has been under Darwinian positive selection. The deletion of CGTG in Tibetan pigs decreased the expression levels of *TMPRSS6* mRNA and protein in the liver. The association of the CGTG deletion with blood physiology dissected a blunted erythropoietic response to high-altitude hypoxia in Tibetan pigs.

To test the signal of selection and identify causal sequence variations, resequencing of the entire genomic region of candidate *TMPRSS6* gene was performed in 40 domestic pigs and 40 wild boars across five altitudes. As a result, 708 SNPs were identified, in addition to one specific CGTG deletion in intron 10 highly prevalent in Tibetan pigs (53.7% and 62.2%). Haplotype analysis of sequence variants in *TMPRSS6* gene among pig populations across continuous altitudes revealed a hitchhiking effect occurring close to CGTG deletion, suggesting that *TMPRSS6* has been under Darwinian positive selection. Semenza and Wang [71] had proved *HIF-1*, consensus DNA binding site was a nuclear factor that was induced by hypoxia and bound to the hypoxia response element (HRE) and contained CGTG as the conserved core sequence [61]. An extensive sample of 838 domestic pigs across 5 continuous altitudes were collected to detect gene frequency of the CGTG deletion. A strong linear correlation (r = 0.959, p = 0.005) between the CGTG deletion frequency and altitude gradient as well as an increased LD with elevated altitude level revealed that the CGTG indel site might be potentially selected by hypoxia.

Seen from genetic effect size, the CGTG indel showed the strong association with hemoglobin levels in HHA Tibetan pigs and its genetic effect on reducing HGB was up to 13.25 g/L. This result also explained the unchanged HGB in HHA Tibetan pigs compared with the MHA Tibetan pigs because of more CGTG deletion in HHA Tibetan pigs. This finding was consistent with Tibetans with whom the major Tibetan alleles at *EGLN1* and *EPAS1* were also associated with lower hemoglobin concentrations, both of which were associated with the HIF pathway [72–75]. Notably, except for HGB, the effects of the CGTG deletion on blood physiology were slightly different between the two HHA and MHA Tibetan pigs in that RBC was a resulting parameter in the former and MCV in the latter. Correspondingly, on one hand, the decreased RBC improved blood viscosity in HHA Tibetan pigs; on the other hand, the smaller MCV improved blood flow resistant and erythrocyte deformability in MHA Tibetan pigs. At this point, the corresponding findings to Tibetans were limited to insighting into hemoglobin concentration for *EPAS1* and *EGLN1* without detection of hemorheologic parameters. In fact, it seemed to be plausible that decreased RBC and smaller MCV, possibly affected by lowered hemoglobin level, were responsible for blunted erythropoiesis and improved blood viscosity as well as erythrocyte deformability.

In order to identify the regulatory effect of CGTG indel, mRNA and protein expression levels of *TMPRSS6* in HHA Tibetan pigs with DD and II genotype were determined by RT-qPCR and western-blot experiments. As expected, the CGTG deletion leaded to the lower expression level of protein and mRNA in *TMPRSS6*, suggesting that *HIF* could not bind to HRE of *TMPRSS6* gene in HHA Tibetan pigs with DD genotype under high-altitude hypoxic environment. The matriptase-2, *TMPRSS6* encoding protein, can be modulated by acute iron deprivation [76, 77]. In rats under acute iron deprivation, matriptase-2 protein levels increased to suppress hepcidin production and increase iron level in the liver [76], suggesting a key role of matriptase-2 in the maintenance of tight systemic iron homeostasis. Furthermore, studies also demonstrated that *TMPRSS6* mRNA expression was upregulated by *HIF* in hypoxia [48, 72, 73]. Here, the CGTG deletion of *TMPRSS6* resulted in the loss of a HRE, which down-regulated *TMPRSS6* expression level and consequently induced the lower HGB and the smaller MCV underlying blunted erythropoiesis and improved blood viscosity and erythrocyte deformability.

## Acknowledgments

The authors thank staff of the animal science and veterinary bureau of Shangri-la County for collections of Tibetan pig samples for this study.

## Author Contributions

Conceived and designed the experiments: XYK, HMM, XG. Performed the experiments: XYK, JHQ, JFY, QJ, XRL, BW, DWY, SXL. Analyzed the data: XYK, XXD. Wrote the paper: XYK, XXD, SLY.

## Supporting information

**S1 Table. Sample of Blood DNA extraction**

**S2 Table. Sample of Blood physiological index Determination**

**S3 Table. Primer pairs information for detecting SNPs in coding region and intron in this study**

**S4 Table. Genotype frequency and allele frequency of the 16 SNPs in continuous altitudes pig populations**

**S5 Table. Associations between CGTG insert/deletion and hematologic parameters in LHA pig populations**

**S6 Table. Associations between CGTG insert/deletion and hematologic parameters in MA pig populations**

**S7 Table. Associations between CGTG insert/deletion and hematologic parameters in LA pig populations**

